# High Precision Detection of Rare Splice Isoforms Using Multiplexed Primer Extension Sequencing

**DOI:** 10.1101/331629

**Authors:** Hansen Xu, Benjamin J. Fair, Zach Dwyer, Michael Gildea, Jeffrey A. Pleiss

## Abstract

Targeted RNA-sequencing aims to focus coverage on areas of interest that are inadequately sampled in standard RNA-sequencing experiments. Here we present a novel approach for targeted RNA-sequencing that uses complex pools of reverse transcription primers to enable sequencing enrichment at user-selected locations across the genome. We demonstrate this approach by targeting hundreds to thousands of pre-mRNA splice junctions, revealing high-precision detection of splice isoforms, including rare pre-mRNA splicing intermediates.

## Main text

RNA sequencing (RNA-seq) has greatly expanded our understanding of the variety of splice isoforms that can be generated within a cell. Identification of the small subset of reads that span exon-exon junctions within transcripts has enabled the unambiguous detection of vast numbers of novel splice isoforms in scores of organisms^1,2^. Yet in spite of the power presented by this approach, the sequencing depth necessary to quantitatively detect many splicing events is significantly higher than most experiments generate. While this limitation of whole-transcriptome profiling has been addressed in part by methods that utilize antisense probes^3,4^ or PCR enrichment^5^ to target sequencing coverage to genomic regions of interest, a deeper understanding of the basic mechanisms by which splicing is regulated, and the pathological consequences of its mis-regulation, will be facilitated by methods that enable detection of splicing states within cells at higher resolution and with greater precision. Towards this end, we have designed and implemented a novel targeted sequencing method that enhances splice junction detection and allows for genome-wide resolution of splicing intermediates. Building upon the historically validated use of primer extension as a tool for assessing splicing status, we demonstrate the ability to multiplex primer extension assays and evaluate the products by deep sequencing, an approach we hereafter refer to as Multiplexed Primer Extension sequencing, or MPE-seq (Fig1A).

**Figure1:**
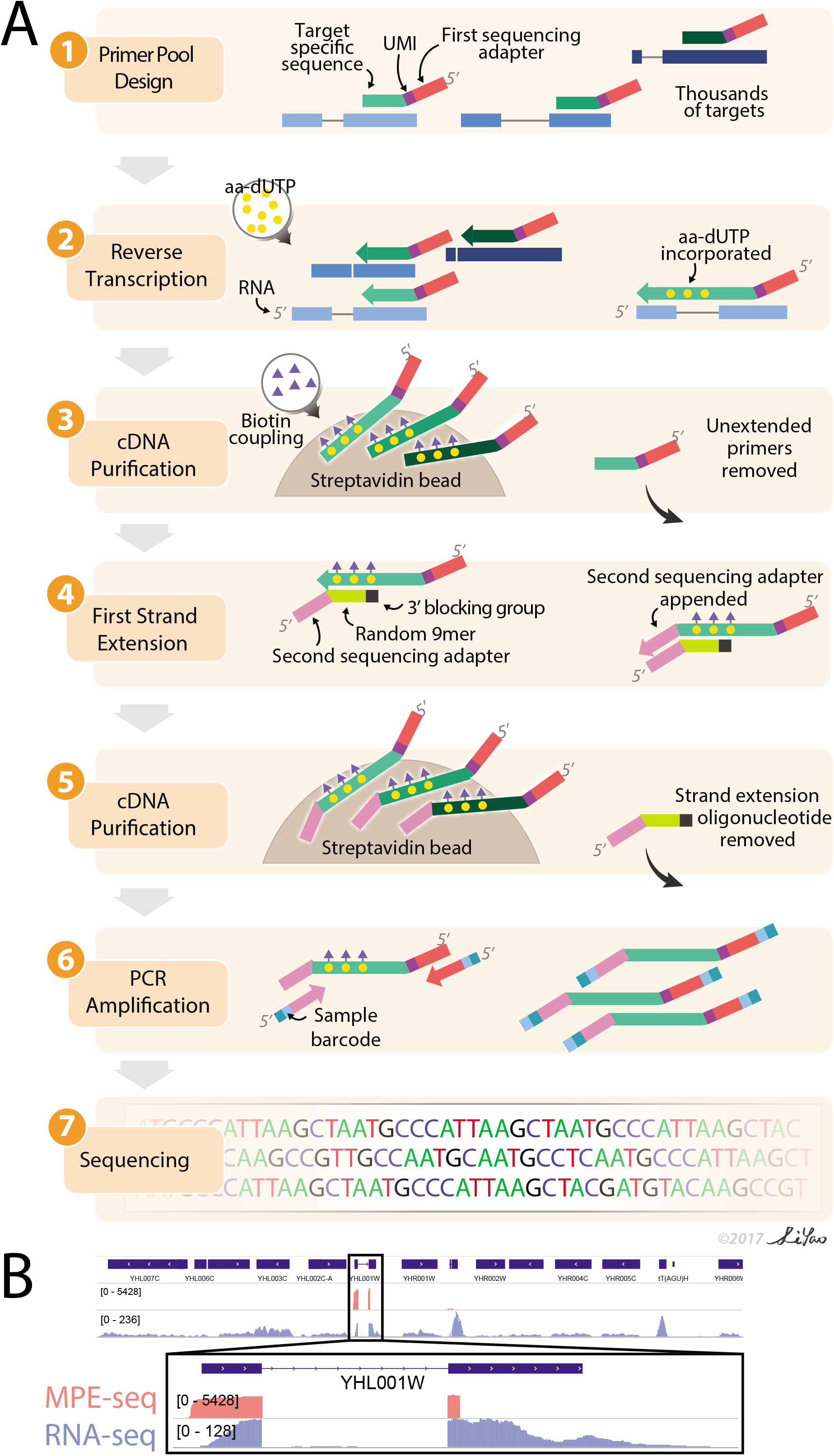
MPE-seq uses complex pools of reverse transcription primers to target sequencing to regions of interest. (A) MPE-seq comprises the following steps: (1) Primers containing sequencing adapter overhangs and a target-specific region are designed for genomic regions of interest. (2) Primers are pooled and used to reverse transcribe total cellular RNA in the presence of amino-allyl dUTP (aa-dUTP), which allows (3) biotin coupling and purification of cDNAs from other reaction components. (4) A second sequencing adapter is appended at the 3’ terminus of the cDNA through first strand extension using Klenow. Libraries are further purified (5), PCR amplified (6) and sequenced (7). (B) Genome browser screenshot of a targeted region in MPE-seq (pink) and conventional RNA-seq (purple)

The method we developed harbors two straightforward yet key features. First, user-selected primers are used to generate complementary DNA (cDNA) during a reverse transcription reaction, enabling targeting of RNA regions of interest. Each primer is appended with a next-generation sequencing adapter, as well as a unique molecular identifier (UMI) to alleviate artifacts associated with PCR amplification during library preparation^6^. The use of elevated temperatures during the reverse transcription reaction minimizes non-specific primer annealing (FigS1), while the inclusion of derivatized nucleotides allows for the efficient purification of extended products and removal of excess primers. Secondly, a strand-extension step similar to template-switching^7^ is used to append the second sequencing adapter onto the 3’ terminus of the cDNA molecules. Coupling this approach with paired-end sequencing allows for the simultaneous querying of the 5’ and 3’ ends of the cDNAs from targeted regions (see Methods for full details).

As an initial demonstration of MPE-seq, we examined pre-mRNA splicing in the budding yeast *Saccharomyces cerevisiae*. For each of the 309 annotated introns in the yeast genome, primers were systematically designed within a 50nt window immediately downstream of the 3’ splice site, ensuring that short extensions would be sufficient to cross the splice junctions. Primers were pooled at equimolar concentration and MPE-seq libraries were generated using total cellular RNA from wildtype yeast and sequenced to a depth of only ~5 million reads. As a comparative reference, standard RNA-seq libraries were generated using poly-A selected RNA and sequenced to ~40 million reads. Whereas the standard RNA-seq libraries yielded read coverage that comprised full gene bodies across the transcriptome, MPE-seq coverage was focused on the selected genes, precisely targeted to the regions upstream of the designed primers (Fig1B). Just over 75% of sequenced fragments from MPE-seq mapped to targeted regions (Fig2A, Supplementallnformation_Table1), resulting on average in a greater than 100-fold enrichment in sequencing depth at these regions when compared with RNA-seq (Fig2B, FigS2). The fold-enrichment was similar across transcripts with a wide range of expression levels (FigS2), and from these data we extrapolate that a standard RNA-seq experiment would require ~500 million sequencing reads to achieve a similar level of coverage over the targeted regions as these 5 million MPE-seq reads provided.

**Figure2:**
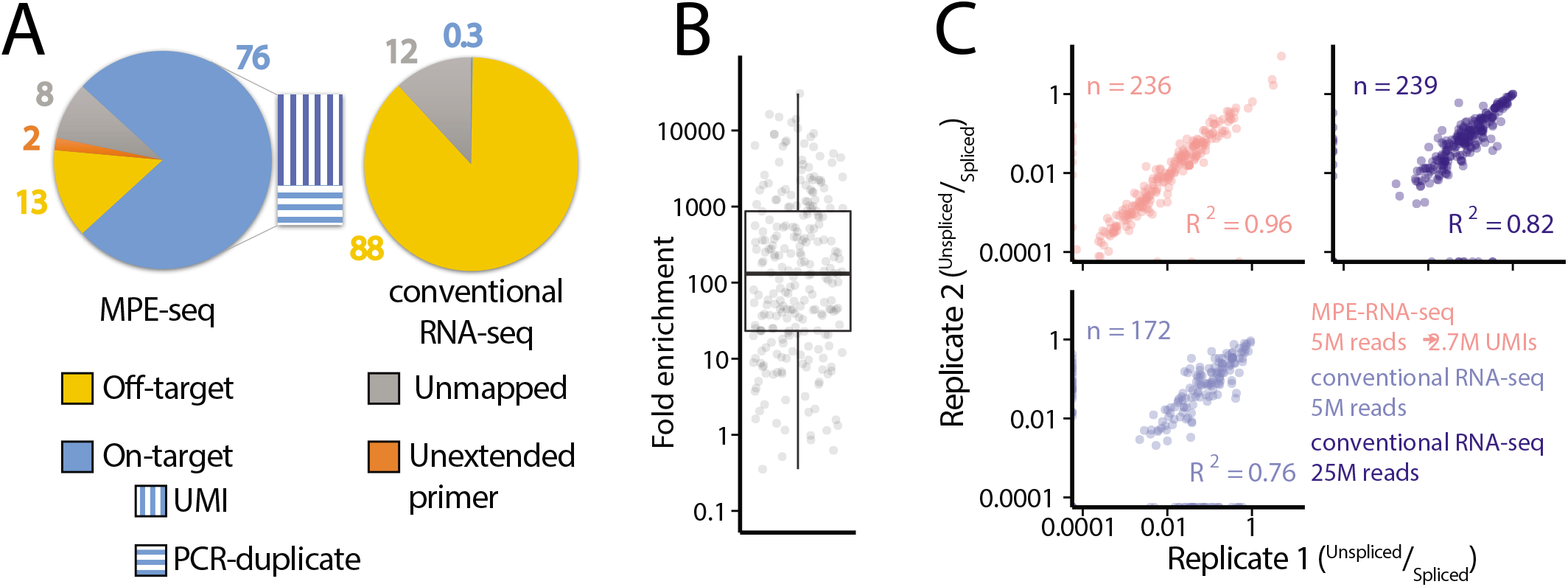
MPE-seq enrichment enables high-precision measurements of splicing. **(A)** The percentage of reads mapped to target and off-target regions is depicted for MPE-seq and conventional RNA-seq. In MPE-seq a small fraction of reads were categorized as “Unextended primer” which corresponds to short primer extension products (0-5 bases extended past the primer) and thus they were not categorized as cDNAs derived from RNA targets. **(B)** Each point represents the fold enrichment of a target region in MPE-seq over conventional RNA-seq. Horizontal lines in boxplots represent the 25^th^, 50^th^, and 75^th^ percentiles. Whiskers end at the 0^th^ and 100^th^ percentiles. **(C)** Scatter plots depict intron-retention measurements in replicate libraries in MPE-seq and conventional RNA-seq at matched or greater read depth. Pearson correlation coefficients (R^2^) are indicated.

Given the increased read depth achieved over targeted regions using MPE-seq, we asked how well rare splice events were sampled. Because each primer extension event corresponded to a single RNA molecule, determining relative isoform expression was simplified because read counts did not need to be adjusted for mapping space as in standard analyses of RNA-seq data (see Methods). Importantly, measurements of the fraction of unspliced message from replicate libraries using MPE-seq showed superior internal reproducibility compared with the larger, replicate RNA-seq libraries (Fig2C), likely reflecting the sampling noise associated with RNA-seq data with reduced sequencing depth over the targeted regions. Moreover, while MPE-seq is not amenable to *de novo* discovery of novel splicing events across the entire genome, it did allow for the identification of scores of rare, previously unannotated splicing events at the targeted regions (SupplementalInformation_Table3), consistent with the significantly increased sensitivity of this approach. Nevertheless, while MPE-seq provided increased sensitivity and reproducibility of splicing measurements, estimates of the fraction unspliced determined from MPE-seq in a wildtype strain only modestly correlated with those determined by RNA-seq (FigS3A, FigS3B). Notably, this correlation improved when comparing how these techniques measured *changes* in splicing between samples assayed by the same methodology (FigS3C), presumably reflecting inherent biases^8^ (in fragmentation, ligation, PCR amplification, library size selection, etc.) present in one or both approaches that are internally well controlled.

With the increased resolution provided by this approach, we sought to determine whether we could detect splicing intermediates using MPE-seq. By identifying the locations of reverse transcription stops, primer extension reactions have historically been used to map a variety of biological features, including among others transcription start sites (TSSs)^9^, and the locations of branch sites within the lariat intermediate species of the pre-mRNA splicing reaction^10,11^ (Fig3A). Because the approach we developed anticipated the possibility of mapping the 3’ ends of the cDNA molecules, we examined the locations of those generated by MPE-seq. As expected, the 3’ ends of many cDNAs accumulated at the TSSs as determined by an orthologous method^12^ (FigS4), indicating that reverse transcription generally proceeded to the 5’ terminus of the RNA. Importantly, we also observed many cDNAs which terminated at or near the annotated branchpoint motifs within introns, with decreased read coverage upstream of the motifs, consistent with the inability of reverse transcriptase to read past the branched adenosine in the lariat intermediate species (Fig3B, Fig3C). This drop in read coverage was not apparent in MPE-seq libraries generated from a strain harboring a conditional mutation in Prp2, an RNA helicase required for catalyzing the 1^st^ step of splicing^13^, corroborating that many of these cDNAs originate from lariat intermediates. We noted that these lariat-intermediate derived cDNAs contain a unique signature of mismatches incorporated by reverse transcriptase at the branched adenosine (FigS5), which may serve as a unique tag for *de novo* identification of branch sites in organisms with less well annotated branch sites^14^.

**Figure3:**
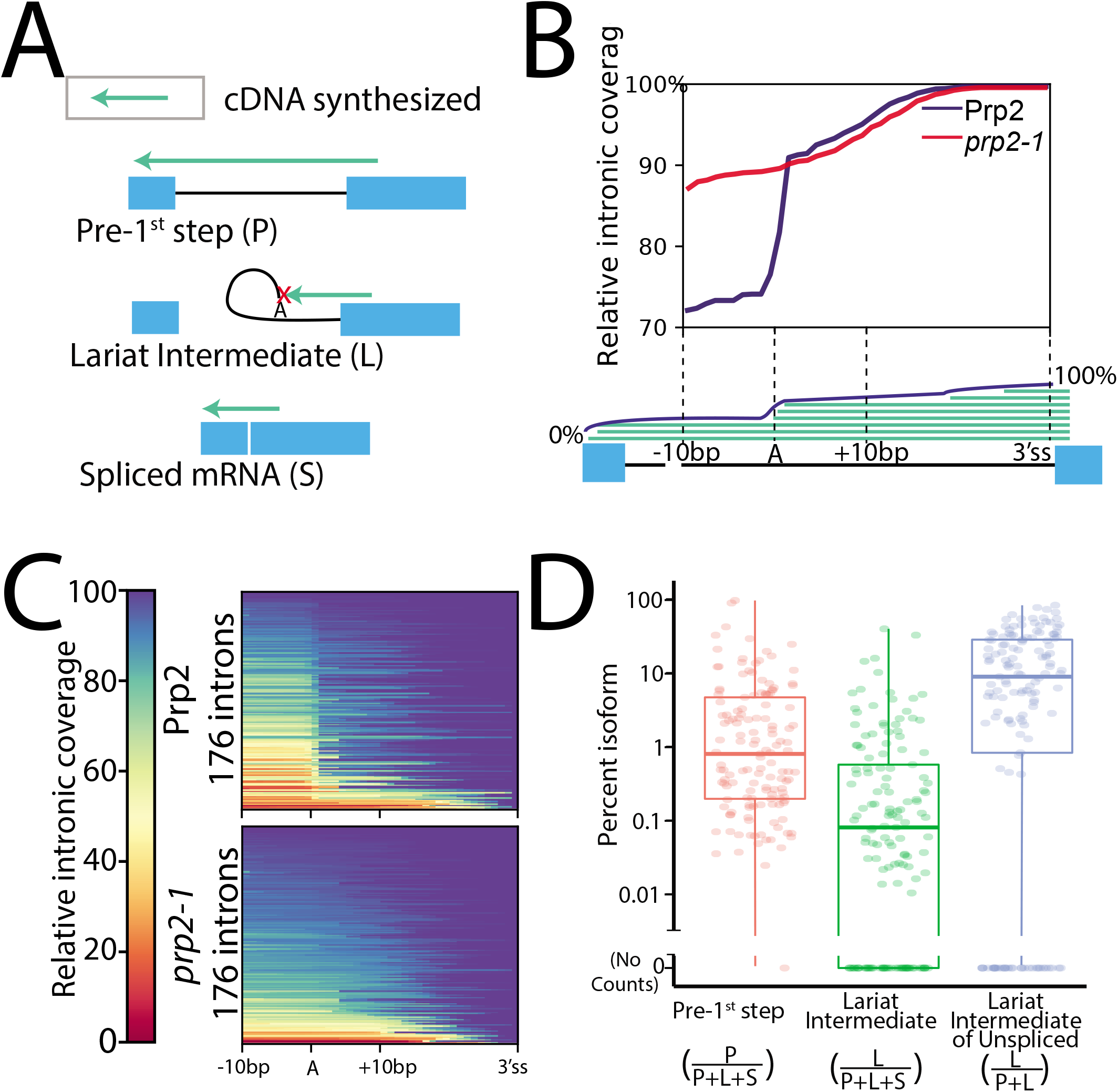
MPE-seq enables genome-wide profiling of lariat intermediates. (A) Schematic depicting cDNA products derived from pre 1^st^ step (P), lariat intermediate (L) and spliced mRNA (S) isoforms. (B) Meta-intron coverage plot surrounding predicted branchpoints in a wild-type (Prp2) and step1 splicing mutant strain (*prp2-1*). The region between the +10 position downstream of the annotated branchpoint and the 3’ splice site (3’ss) was re-scaled for each intron. (C) Heatmap plots showing the relative coverage at each intron for which lariat intermediate reads were detected. (D) Estimates of the relative abundance of each isoform for each targeted intron for which reads were detected. Horizontal lines in boxplots represent the 25^th^, 50^th^, and 75^th^ percentiles. Whiskers end at the 0^th^ and 100^th^ percentiles.

The ability of MPE-seq to differentiate between unspliced isoforms in unfractionated cellular RNA provided a unique opportunity to investigate the relative efficiencies with which transcripts undergo the 1^st^ and 2^nd^ chemical steps in the splicing pathway. Our data revealed that ~10% of unspliced pre-mRNAs at steady state conditions were present in the lariat intermediate form genome-wide (see FigS6, methods), albeit with significant variation between individual pre-mRNAs (Fig3D). Remarkably, a strong correlation was observed between the relative levels of pre-mRNA and lariat intermediate species for a given transcript (FigS7A). Correlations were also observed between splice intermediate levels and transcript expression level, branch motif strength, and host gene function (FigS7B, FigS7C); however, no correlation was seen when considering the ratio of pre-1^st^ step intermediate to lariat intermediate, a metric which we expect to reflect the relative catalysis rates of the 1^st^ and 2^nd^ step of splicing. A complete understanding of the determinants of *in vivo* splicing efficiency will require measurements of the kinetics of these individual steps rather than their steady state levels. The ability of MPE-seq to robustly differentiate pre-mRNA isoforms provides a powerful new opportunity to do just this.

Whereas our initial experiments were performed using individually synthesized oligonucleotides as primers, we sought to increase the utility of this approach by examining methods that would facilitate an increase in the number of targeted regions. Many commercial sources exist that allow for the cost-effective, array-based synthesis of pools of thousands of individual oligonucleotide sequences, so we developed an approach that would enable the use of such primers. A common sequence was appended onto the 3’ end of the desired primers, facilitating a protocol that included PCR amplification, restriction digestion, and targeted strand degradation, to produce single-stranded oligonucleotides with which MPE-seq libraries could be generated (Fig4A, FigS8). Using this approach, we designed oligos that allowed us to re-create the 309 previously described *S. cerevisiae* primers, and an additional 3918 primers that targeted splice junctions in the relatively intron-rich fission yeast *Schizosaccharomyces pombe*. Importantly, genome-wide splicing efficiencies determined from MPE-seq libraries generated using primers from pooled synthesis were highly correlated with splice efficiencies derived from individually synthesized oligos (Fig4B), validating the utility of this approach for oligo synthesis. Moreover, MPE-seq libraries generated with primers derived from pooled synthesis also showed strong enrichment for the targeted regions, with levels on par with what was observed using individually synthesized oligonucleotide primers (Fig 2A and 4C). Not surprisingly, a modest increase in off-target reads was seen when primers from the pooled synthesis were used, consistent with the decreased sequence fidelity of array-based oligo synthesis^15^ and the increased capacity of these aberrant oligos to prime reverse transcription at undesirable locations. Nevertheless, a median enrichment at targeted regions of six-fold was achieved even when thousands of primers were used in fission yeast (TableS9), again enabling detection of rare but natural alternative splicing events^16^ that are poorly sampled using standard RNA-seq library preparation methods (Fig4D), and further confirming the capacity of MPE-seq to effectively target thousands of unique user-selected loci.

**Figure 4:**
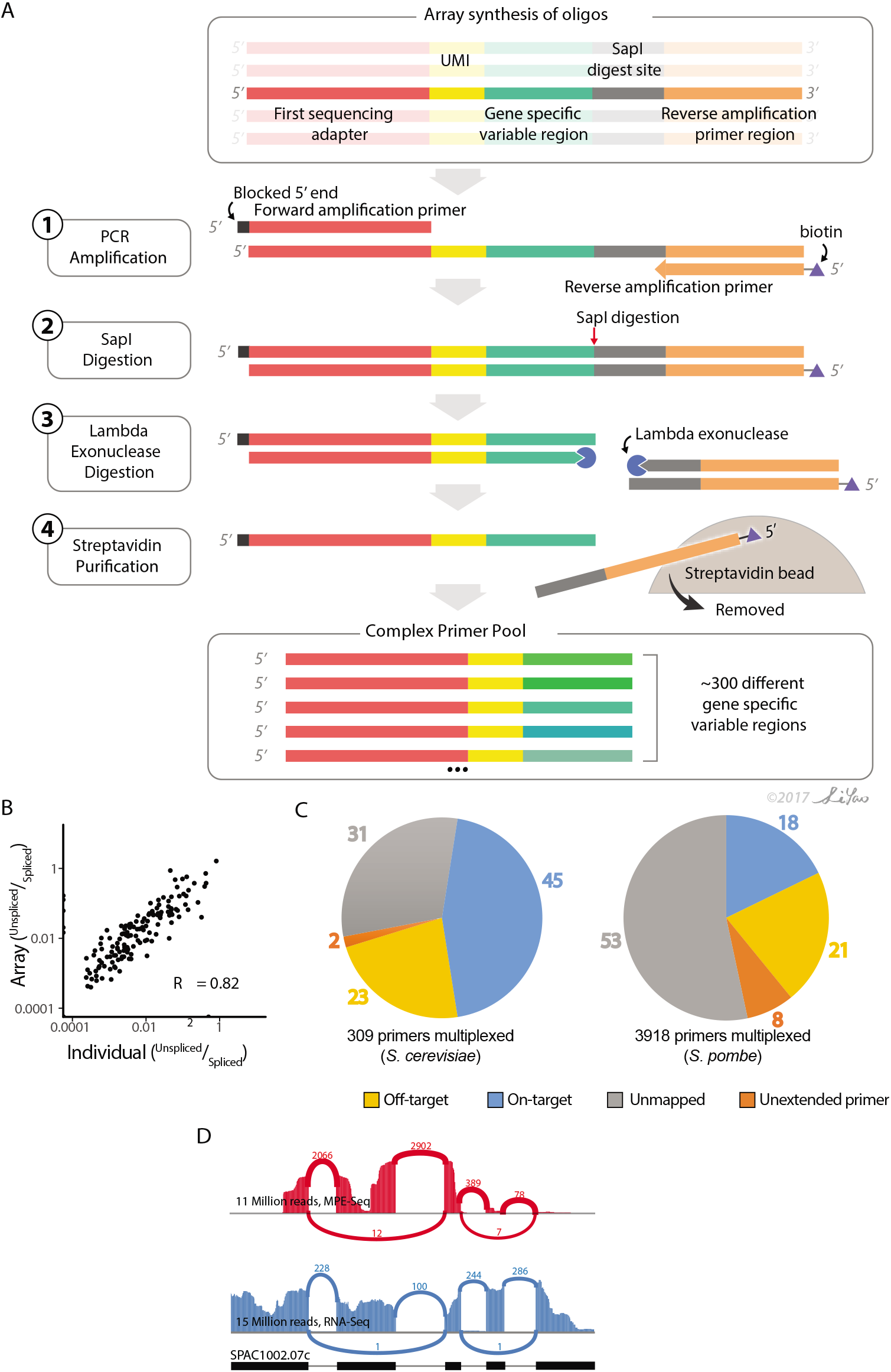
Array-based oligonucleotide synthesis can be used to generate primer pools for use in MPE-seq. (A) Obtaining adequate amounts of primer pools for MPE-seq from cost-effective array-based oligonucleotide synthesis can be achieved in four steps. (1) PCR amplification of the oligonucleotide synthesis pool using a 5’ blocked sense primer and a biotinylated antisense primer. (2) Restriction digestion to cleave off the PCR primer handle. (3) Lambda exonuclease digestion of free 5’ ends. (4) Streptavidin purification of biotinylated PCR handle. The unbound fraction is the desired primer pool product. **(B)** A scatter plot compares the fraction of unspliced mRNA measured by MPE-seq libraries which used individually synthesized primer pools versus array-based synthesis of primer pools. (C) The percentage of reads mapped to target and off-target regions is depicted for MPE-seq using array-synthesized primers. (D) Sashimi plot of a targeted region within the *ats1* gene locus demonstrates the capacity of MPE-seq to reveal complex alternative splicing patterns with higher sensitivity than RNA-seq, despite having lower total sequencing depth.

Our work here demonstrates the capacity of MPE-seq to facilitate examinations of pre-mRNA splicing status in a targeted, cost-effective way that improves the precision and sensitivity of splice isoform detection. The improved sensitivity of this approach is perhaps best exemplified by our ability to detect the lariat intermediate products of the pre-mRNA splicing pathway. Though other recently described methods^17^, including splicoesome profiling^14,18^, have reported large-scale detection of upstream exon splice intermediates and excised lariats, MPE-seq uniquely detects lariat intermediates, not excised lariats, from unfractionated cellular RNA. Moreover, the ‘profiling’ methods, which detect RNAs physically associated with the splicoesome, require protein tagging and/or purification steps that require large amounts of starting material, limiting the types of applications to which they might be applied. Conversely, MPE-seq can be implemented in virtually any system of interest with a need for only microgram quantities of RNA. Additionally, the ability of MPE-seq to query RNA from a wide variety of sources (cytoplasmic/nuclear fractionated RNA, polysome-fractionated RNA, poly-A selected RNA, metabolically labelled RNA) enables an analysis of the cellular location, translational or polyadenylation status and turnover rates of splice isoforms and intermediates.

Importantly, our demonstration that oligonucleotides derived from pooled commercial syntheses can be used in MPE-seq expands the types of applications to which this approach could be applied in a cost-effective manner. While we see no *de facto* limitation to the number of unique primer sequences that could be used for MPE-seq, with increasing numbers of primers comes an increasing potential for their cross-reactivity with undesirable RNA targets, highlighting the importance of specificity and fidelity in primer design and synthesis. Similarly, the level of enrichment provided by this approach will vary as a function not only of the number of regions being targeted, but also of the distribution of the expression levels of those targets. Whereas standard RNA-seq experiments suffer from over-sampling of highly expressed transcripts at the expense of under-sampling lowly expressed transcripts, targeted approaches like MPE-seq offer the opportunity to reduce this problem by grouping targets with similar expression levels, enabling similar coverage over a range of expression levels. Overall, we expect that the improved sensitivity, precision and flexibility of this approach will enable a higher-resolution understanding of the pre-mRNA splicing pathway. Likewise, primer extension assays have been used to assay RNA secondary structure after *in vitro*^19^ or *in vivo*^20^ chemical probing, and we expect that MPE-seq could be readily adapted to RNA structure interrogation and other approaches where primer-extension assays or targeted RNA sequencing is applicable.

## Online Methods

### Strain Maintenance and Growth Conditions

Unless otherwise indicated, all *S. cerevisiae* experiments used the wild type (WT) strain BY4741 (*MATa, his2Δ1, leu2Δ0, met15Δ0, ura3Δ0*). Single colonies were inoculated into liquid YPD media and grown overnight at 30□. Overnight cultures were then inoculated into fresh liquid YPD, seeding cultures at OD_600_ ~0.05. Cells were collected by vacuum filtration once cultures reached OD_600_ ~0.7, immediately followed by flash freezing in liquid nitrogen. Cell pellets were then stored at −80□. For the temperature sensitive mutant *prp2-1*^21^ we grew cultures as described above except at 25□. Once cultures reached OD_600_ ~0.7, an equal volume of fresh 50□ YPD media was added to shift cells to the non-permissive temperature of 37□. The cultures were then maintained at 37□ for 15 minutes before cell collection as described above. All *S. pombe* experiments used the wild type strain JP002 (*h*^+^, *ade6-M210, leu1-32, ura4-D18*). Single colonies were inoculated into liquid YES media and grown overnight at 30°C. Overnight cultures were then inoculated into fresh liquid YES, seeded at OD_600_ ~0.05. Cells were collected by vacuum filtration upon reaching OD_600_ ~0.5 and immediately flash frozen in liquid nitrogen. Cell pellets were stored at −80°C.

## MPE-seq Primer and Oligo Design

### Gene specific reverse transcription primer design

For each of the 309 annotated spliceosomal introns within the *S. cerevisiae* genome (annotations obtained from UCSC SacCer3) as well as for a subset of introns (3918 in total) within the *S. pombe* genome (annotations obtained from Ensemble ASM294v2.37), a reverse transcription primer was designed within the first 50 nucleotides downstream of the intron. Targeting to this region ensured that short-read sequencing of the products generated from reverse transcription with these primers would cross the upstream exon-exon or exon-intron boundaries, enabling determination of the splicing status. Primers were designed using OligoWiz, a program initially developed for microarray probe design, but which enables the selection of primer sequences optimized for target specificity relative to a designated genomic background^22^. We used the standalone version of OligoWiz with default parameters for short (24-26bp) oligo design to obtain optimal sequences within each 50bp window. To the 5’ end of each of these sequences was appended two additional sequence elements: a random 7-nucleotide unique molecular index (UMI) which allows for the detection and removal of amplification artifacts arising from library preparation^6^; and the P5 region of the Illumina sequencing primer to enable the sequencing of the reverse transcription products. Each of the primers targeting *S. cerevisiae* junctions was individually synthesized by Integrated DNA Technologies (IDT), the full sequences of which are provided in (Supplemental Information Table 5). Array-based oligonucleotide synthesis was performed by LC Sciences using individual OligoMix syntheses for primers from each species (Supplemental Information Table 8).

### Complex oligo mix amplification method

The array-based oligos are synthesized at vastly lower quantities than is required for cDNA synthesis in MPE-seq. To generate a quantity of primer pool that is sufficiently large, PCR amplification along with several processing steps were used (Fig4A). This was enabled by addition of 2 key sequence elements appended onto the 3’ end of the individually synthesized oligo primers detailed above. From the 5’ to 3’ direction: (1) a SapI restriction site; and (2) a PCR amplification sequence (Supplemental information Table 5). The oligos were amplified in a standard PCR reaction using Phusion polymerase. This 400 μL PCR reaction contained: 1% of the pooled oligonucleotides from LCSciences as a template, a forward primer (oHX093) containing a C3 spacer at its 5’ end, and a reverse amplification primer (oHX094) containing a biotin-label at its 5’ end (see Supplemental Information Table 6). A total of 14 amplification cycles were performed, each consisting of the following conditions: denaturation at 95°C for 10 sec; annealing at 60°C for 20 sec; and extension at 72°C for 30 sec. Upon completion of this initial reaction, the entire reaction was used as a template to seed a larger (40 mL) PCR reaction. For efficient amplification, this large reaction was performed in four 96-well plates with 100μL in each well. Reaction conditions were identical to those described for the first reaction, but a total of 15 cycles were performed for this second amplification. Reactions were purified and concentrated by isopropanol precipitation. To generate single stranded primers for use in MPE-seq, the double-stranded amplicons were first digested using SapI (NEB R0569) in a 150 μL reaction containing 30 μL of enzyme. The reaction was incubated at 37°C overnight, after which the reaction products were concentrated by ethanol precipitation. Next, the 5’ to 3’ lambda exonuclease (NEB M0262) was used to preferentially degrade the two strands containing unmodified 5’ ends. This reaction was performed at 37°C for 2 hours according to the manufacturer’s protocol. The products of this reaction were then purified using Zymo columns using 7X volume binding buffer (2 M guanidinium-HCl, 75% isopropanol). After this step, the remaining DNA consisted of the desired single stranded RT primer, and an undesired single stranded section containing the SapI site plus the amplification primer. Making use of the 5’ biotin tag on the amplification primer, these undesired oligos were removed by affinity capture with streptavidin beads. Specifically, this was accomplished by using 50 μL of Dynabeads MyOne Streptavidin C1 according to the manufacturer’s protocol. The unbound supernatant fraction was retained as it contains the desired products. The recovered material was precipitated and verified using 6% native PAGE stained with SyBr Gold.

### 1^st^ Strand extension template oligo design

The oligos were designed with 3 key features from the 5’ to 3’ end of the oligo. First, a portion of the Nextera P7 sequencing adapter. Of the entirety of the P7 adapter, the region 3’ of the i7 barcode was used. This allowed for barcoding and amplification of the sequencing libraries. Second, a dN9 or dN12 anchor on the 3’ end of the oligo allowed it to randomly anneal to cDNA products. And third, a 3’ carbon block modification (hexanediol, IDT) was added to preclude the ability of Klenow to extend this primer. As such, the oligo may only be used as a template to append the Nextera sequencing adapter onto the end of first strand cDNAs. The full sequence of this primer can be found in (Supplemental Information Table 6).

## MPE-Seq Library Prep

cDNA synthesis. For *S. cerevisiae* libraries, RNA was isolated following a hot acid phenol extraction protocol^23^. Each library was generated using 10 μg of total RNA. From this RNA, cDNA was synthesized by mixing 1 μg of the gene specific primer pool described above with each RNA sample in a 20 μL reaction containing 50 mM Tris-HCl (pH 8.5), 75 mM KCl. The primers were then annealed in a thermocycler with the following cycle: 70°C for 1 minute; 65°C for 5 minutes; hold at 47°C. An equivalent volume of MMLV reverse transcriptase enzyme mix containing 1 mM dATP, 1 mM dGTP, 1 mM dCTP, 0.4 mM aminoallyl-dUTP, 0.6 mM dTTP, 50 mM Tris-HCl (pH 8.5), 150 mM KCl, 6 mM MgCl2, 10 mM DTT was pre-heated to 47°C and added to the primer-annealed RNA mix, resulting in a total reaction volume of 40 μL. Maintaining the samples at 47°C was essential for reducing off-target cDNA synthesis. Reactions were incubated at 47°C for 3 hours, followed by heat inactivation at 85°C for 5 minutes. Remaining RNA was hydrolyzed by addition of ½ volume of 0.3 M NaOH, 0.03 M EDTA and incubation at 65°C for 15 minutes. After neutralization with ½ (original) volume of 0.3 M HCl, the cDNA was purified with a Zymo-5 column using 7X volume of binding buffer (2 M guanidinium HCl, 75% isopropanol). Purified cDNA samples were dried to completion in a SpeedVac. For *S. pombe* libraries, RNA was isolated as described above. Poly-adenylated RNA was then isolated from 60 μg of total RNA using the NEBNext Poly(A) mRNA Magnetic Isolation Module. RNA was then fragmented to an average size of 200 nucleotides by incubating in 10 mM ZnCl_2_, 10 mM Tris-HCl pH 7.0 for 10 minutes at 65°C. The reaction was then quenched by addition of EGTA (pH 8.0) to a final concentration of 50 mM. The cDNA synthesis reactions were performed as above with some modifications. For reasons described below, 4 μL of Superscript III (ThermoFisher) reverse transcriptase was used along with the manufacturer supplied 5x buffer. For primer annealing and extension samples were held at 55°C for 1 hour, followed by heat inactivation at 85°C for 5 minutes.

### NHS ester biotin coupling

Dried cDNA samples were resuspended in 18 μL of fresh 0.1 M Sodium Bicarbonate (pH 9.0), to which 2 μL of 0.1 mg/μL NHS-biotin (ThermoFisher 20217) was added. Reactions were incubated at 65°C for 1 hour followed by purification of biotin coupled cDNA from unreacted NHS-biotin by using Zymo-5 columns using 7X volume of binding buffer (2 M guanidinium HCl, 75% isopropanol).

### Streptavidin-biotin purification

20 μL of Dynabeads MyOne Streptavidin C1 (ThermoFisher 65602) per sample were pre-washed twice in 500 μL of 1X bind and wash buffer (5 mM Tris-HCl (pH 7.5), 0.5 mM EDTA and 1 M NaCl) as per manufacturer’s protocol. Washed beads were resuspended in 50 μL of 2X bind and wash buffer per sample and 50 μL was combined with each 50 μL purified cDNA sample. Biotin-streptavidin binding was allowed to proceed for 30 minutes at room temperature with rotation. Bound material was washed twice with 500 μL of 1X bind and wash buffer, followed by an additional wash with 100 μL of 1X SSC. To ensure purification of only single-stranded cDNAs, beads were then incubated with 0.1 M NaOH for two consecutive room temperature washes for 10 minutes and 1 minute, respectively. Finally, the bound material was washed 3 times with 100 μL 1X TE. The cDNA was eluted from the beads by heating samples to 90°C for 2 minutes in the presence of 100 μL of 95% formamide, 10 mM EDTA. The eluate was then purified using Zymo-5 columns as described above, and the cDNA was eluted from columns in 40 μL of water.

### First strand extension

Primers were annealed to purified cDNA by combining: 1uL 1^st^ strand extension oligo (100 μM oJP788 for *S. cerevisiae* and oJP789 for *S. pombe*), 5 μL 10x NEB buffer 2, 40 μL purified cDNA sample, and 1 μL of 10 mM (each) dNTP mix. Samples were then incubated at 65°C for 5 minutes, followed by cooling to room temperature on the bench top. To each sample was added 3 μL of Klenow exo- fragment (NEB M0212) and reactions were incubated for 5 minutes at room temperature, after which they were moved to 37°C for 30 minutes. Samples were subsequently purified using streptavidin beads following the protocol described above. Samples were then concentrated using Zymo-5 columns, with elution in 33 μL of water.

### PCR amplification

Amplification of the reaction products was accomplished by using 10 μL of the purified material generated in the 1^st^ strand extension reaction as a template in a PCR reaction. Illumina Nextera (i5) and (i7) indexing primers were used in a standard 50 μL PCR reaction with Phusion polymerase (ThermoFisher F530S). Cycling conditions were as follows: denaturation at 95°C for 10 sec; annealing at 62°C for 20 sec; and extension at 72°C for 30 sec. Libraries typically required between 14 and 20 cycles of amplification, depending upon the efficiency of library preparation. Each PCR reaction was then run on a 6% native poly-acrylamide gel, and the DNA was resolved by staining with SyBr gold. Libraries were size selected from 200bp to 800bp and DNA was extracted from gel fragments via passive diffusion overnight in 0.3 M sodium acetate (pH 5.3). Libraries were then ethanol precipitated and quantified.

### cDNA synthesis temperature experiment

Due to the target specific nature of MPE-seq cDNA synthesis, any reverse transcription (RT) events at non-target sites will reduce the fraction of on-target reads. Indeed, these off-target events contribute significantly to the nonspecific class reads in a typical MPE-seq experiment (Fig2A). One way to reduce off-target RT events is through increasing the specificity of the RT primers. We assessed this by testing the effect of increased temperature during the RT reaction on off-target sequencing reads. MPE-seq libraries were generated using the above described protocol with one primary difference: Increased reaction temperatures required the use of a thermostable enzyme. For this reason, Superscript III (ThermoFisher) was used along with the manufacturer supplied buffer (reaction concentration: 50 mM Tris-HCl pH 8.3, 75 mM KCl, 3 mM MgCl2). Primer annealing and reactions were carried out at 47°C, 51°C, and 55°C in replicate.

## MPE-Seq Data Anlysis

### Sequencing and alignment

*S. cerevisiae* MPE-seq libraries were sequenced on the NextSeq platform by the BRC Genomics Facility at Cornell university using 60bp (P5) + 15bp (P7) paired-end chemistry. PCR duplicates were removed from the dataset by filtering out non-unique reads with respect to all base calls in both reads, including the 7bp UMI. In other words, for each set of identical paired-end reads, a single read-pair was retained for analysis. MPE-seq reads were aligned to the yeast genome (Reference genome assembly R64-1-1^24^) using the STAR aligner^25^ with the following alignment parameters: {--alignEndsType EndToEnd -- alignIntronMin 20 --alignIntronMax 1000 --alignMatesGapMax 400 -- alignSplicedMateMapLmin 16 --alignSJDBoverhangMin 1 --outSAMmultNmax 1 -- outFilterMismatchNmax 3 --clip3pAdapterSeq

CTGTCTCTTATACACATCTCCGAGCCCACGAGAC --clip5pNbases 7 0}. Alignment files were filtered to exclude read mappings deriving from inserts less than 30bases. We believe these short fragments represent unextended reverse transcription primers that were retained in the sequencing libraries. These small fragments can sometimes erroneously map to splice junctions or target introns, even though we believe they are not derived from cellular RNA. *S. pombe* MPE-seq libraries were sequenced on the MiSeq platform by the BRC Genomics Facility at Cornell university using 100bp (P5) + 50bp (P7) paired-end chemistry. Reads were trimmed to 60bp and 15bp and processed as above for Fig4C while full length reads were processed as above for Fig4D.

*S. cerevisiae* RNA-seq libraries were sequenced on an Illumina HiSeq2500 by the BRC Genomics Facility at Cornell University using 100bp single end reads. *S. pombe* RNA-seq data^26^ were downloaded from NCBI (accession code SRS167019) and R2 of read pairs was discarded to make read lengths comparable to our other libraries. Reads were aligned using the STAR aligner with the following alignment parameters: {--alignEndsType EndToEnd --alignIntronMin 20 --alignIntronMax 1000 --alignSJDBoverhangMin 1 --outSAMmultNmax 1 -- outFilterMismatchNmax 3 --clip3pAdapterSeq CTGTCTCTTATACACATCTCCGAGCCCACGAGAC}.

When applicable, replicate libraries were combined prior to alignment. However, to assess technical reproducibility of MPE-seq, replicate libraries were subsampled to varying read depths, aligned separately, and compared to RNA-seq libraries also subsampled to varying read depths. All data are available through NCBI’s GEO at accession number XXXX.

## Estimating fraction of on-target reads and MPE-seq enrichment

Using bedtools^27^, read 1 alignments extending into a targeted intron or crossing a targeted exon-exon junction were considered on-target. Read 1 alignments which mapped downstream of a targeted intron but did not extend into an intron or cross an exon-exon junction were considered unextended primers. Read 1 alignments that mapped to the genome at non-targeted loci were considered off-target. Unmapped reads were then realigned to the genome with the same parameters as above (except with --clip5pNbases 31 0) and subsequent read 1 alignments to non-targeted loci were also considered off-target. Remaining reads were considered unmapped. Enrichment was calculated by dividing the number of read-count normalized exon-exon junctions found in MPE-seq by the number of read-count normalized exon-exon junctions in RNA-seq datasets for each targeted intron.

## Estimating splice isoform abundances from MPE-Seq data

For each intron, the relative abundance of unspliced and spliced isoforms was determined by counting spliced and unspliced reads. Spliced reads (*S*) were counted using the SJ.out.tab file created by the aligner. Unspliced reads were counted using bedtools^27^ to count the number of reads that cover any part of the intron, considering only the first read of the paired-end reads. Unspliced read counts were further categorized as deriving from a lariat intermediate (*L*) or pre-1st step RNA (*P*) by considering the mapping location of the second read of the paired end reads, which we observed to often terminate near the TSS, or in the case of a lariat-intermediate-derived cDNA, near the branchpoint-A of the intron. Based on paired end mapping locations, each fragment was categorized into one of six categories (See FigS6) and the counts within those six categories were used to calculate *S, P*, and *L* as follows:

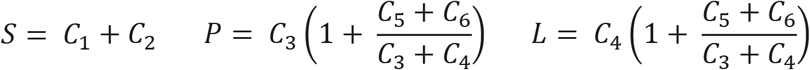

Locations of branchpoints (Supplementary Information Table 7) were determined by consolidating the most used branchpoint from lariat sequencing data^28^ and previously described branch locations based on sequence motif searches^29^.

### Heatmaps and meta-gene plots

To generate metagene plots which illustrate read coverage around features of interest, we used the deepTools ComputeMatrix command^30^ in conjunction with a BigWig coverage file of the 3’ terminating bases and a bedfile containing TSS-positions as determined by PRO-cap^12^ or a bedfile containing the annotated branchpoint regions detailed above. Importantly, this bedfile was filtered to only include branchpoint regions that would produce a lariat intermediate that is within the size range captured by library size-selection of MPE-seq libraries (see column “AttemptedLariatQuantification?” in Supplemental Information Table 2).

## RNA-seq Experiments

### Library prep

For each RNA-seq library, 1 μg of total RNA was input into the “NEBNext Ultra Directional RNA Library Prep Kit for Illumina”. Libraries were prepared following the manufacturer’s protocol.

### Estimating Splice isoform abundances from RNA-seq data

Similar to MPE-seq data, spliced reads from target introns were counted using the SJ.out.tab file created by the aligner. Unspliced reads were counted using the bedtools software package^27^ to count the number of reads which overlapped an intron. Spliced and unspliced read counts for each intron were then length normalized for the feature’s potential mapping space. The potential mapping space for a spliced read is equal to 2 x read length minus the minimum splice junction overhang length. The potential mapping space for an unspliced read is equal to the 2x read length minus minimum splice junction overhang length plus length of the intron. Reads counts assigned to each feature were then divided by the length. Fraction unspliced was calculated for each intron as the quotient of length normalized unspliced reads and spliced reads. Relative transcript expression was calculated via transcripts per million (TPM) normalization^31^, only considering exonic reads and exonic gene-lengths.

## Acknowledgements

We thank members of the J.A.P., H. Kwak, and A. Grimson laboratories for critical feedback on this work. We thank Li Yao for initial drafts of Fig1A and Fig4A. We thank P. Schweitzer and the BRC Genomics Facility at Cornell for outstanding technical support with Illumina sequencing. This work was funded by a Research Scholars Grant from the American Cancer Society (to J.A.P.) and NIH grant R01GM098634 (to J.A.P.)

## Author Contributions

All authors contributed to research design. H.X., B.F., Z.D. and M.G. performed research and analyzed data. All authors wrote the paper.

---

**FigureS1:**
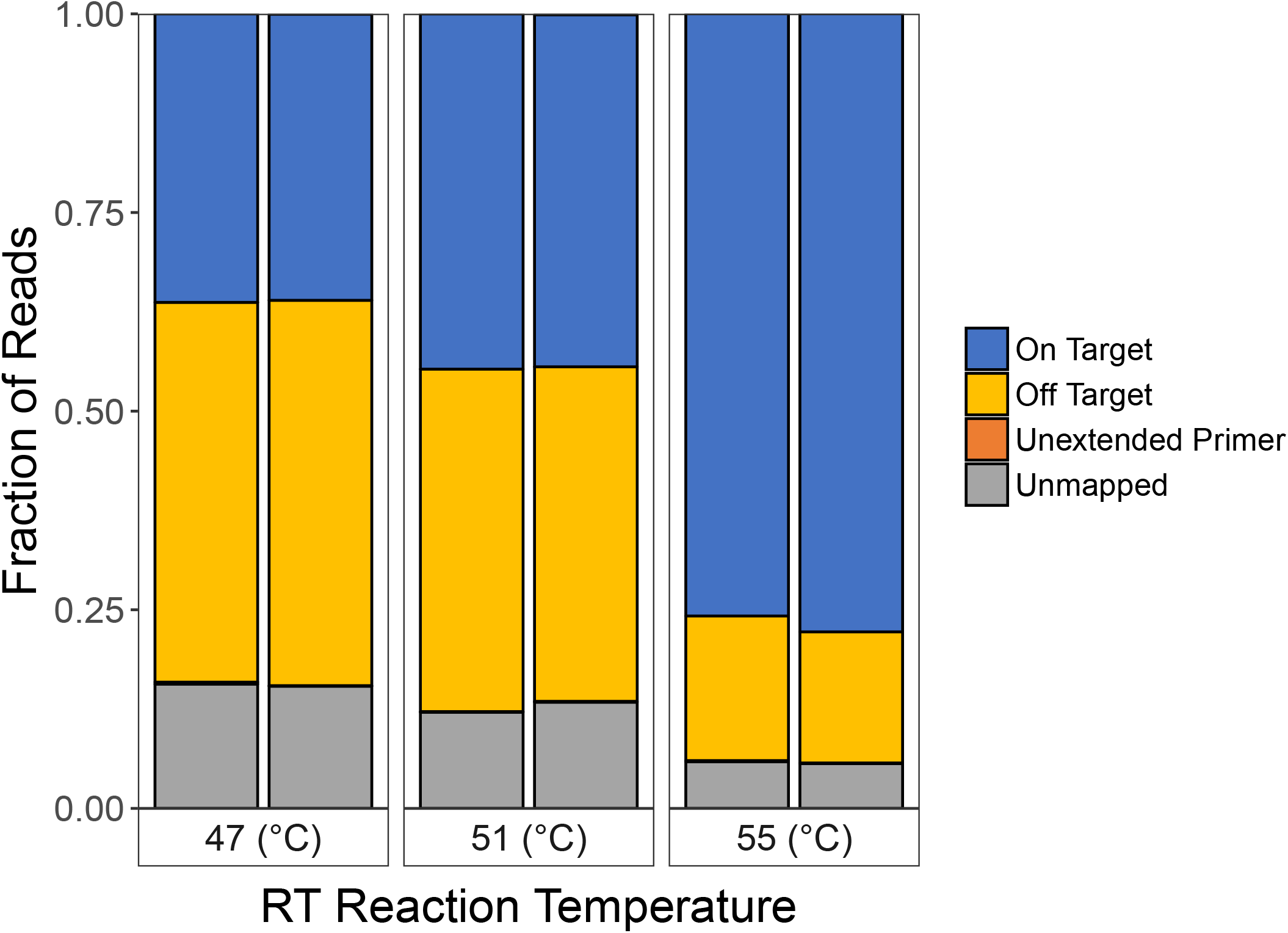
Elevated temperatures in reverse transcription reactions increase specificity. The fraction of on-target and off-target reads from replicate MPE-seq libraries generated from reverse transcription reactions performed at various temperatures. A small fraction of reads were categorized as “Unextended primer” which corresponds to short primer extension products (0-5 bases extended past the primer) and thus they were neither categorized as cDNAs derived from RNA targets or unamappable.

**FigS2:**
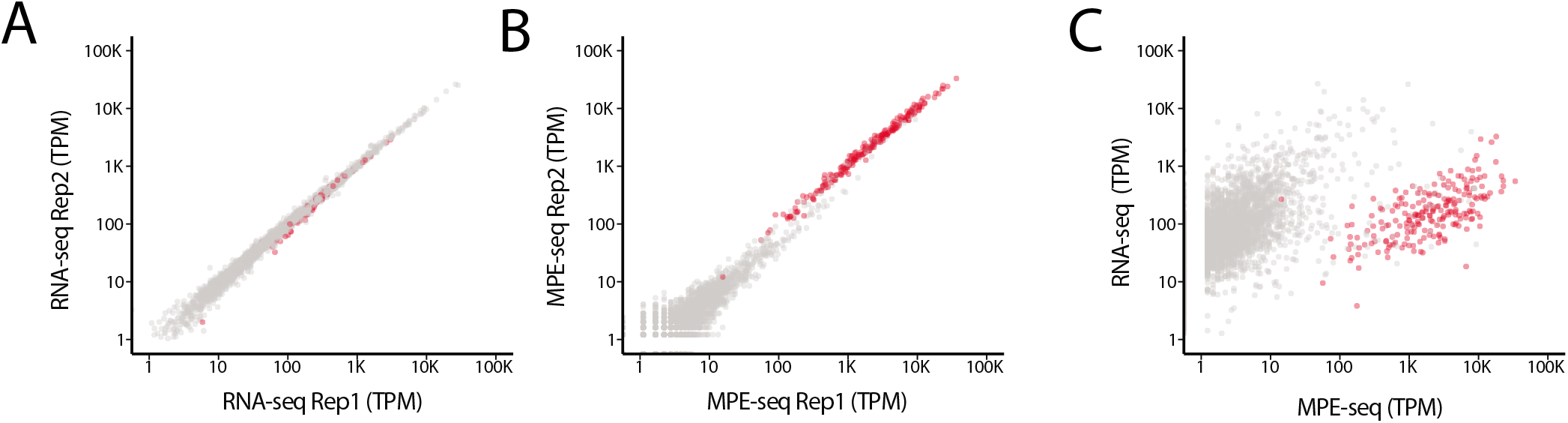
Expression measurements as determined by MPE-seq and RNA-seq. **(A)** A scatter plot depicts gene expression measurements (RNA-seq) in replicate datasets. Genes containing splice-events that were among those chosen for targeted sequencing are depicted in red. These targeted genes range in expression levels by orders of magnitude. **(B)** A scatter plot depicts gene expression measurements in replicate MPE-seq datasets (red). Similar to conventional RNA-seq, expression measurements in MPE-seq are highly reproducible between replicates, even for the small proportion of mis-priming events that map to off-target locations (grey). **(C)** A scatter plot depicts gene expression measurements in RNA-seq and MPE-seq. The right shift of targeted genes reflects successful enrichment of targets by orders of magnitude. The observation that even highly expressed genes as measured by RNA-seq are proportionally highly expressed in MPE-seq suggests that primers are not limiting during reverse transcription.

**FigS3:**
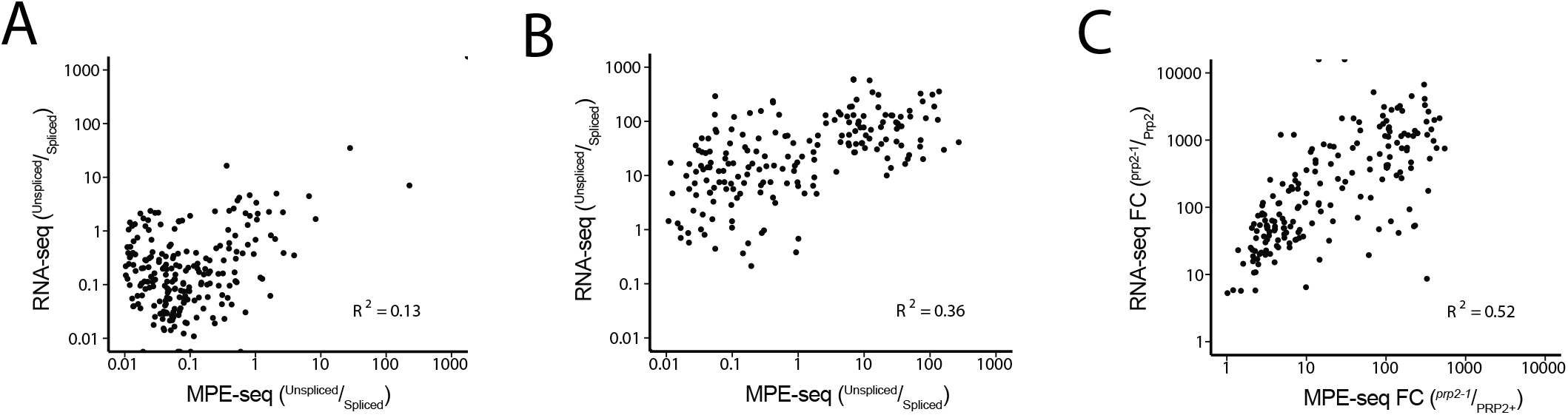
Splicing measurements as determined by MPE-seq and RNA-seq. **(A)** A scatter plot depicts intron-retention measurements in MPE-seq and conventional RNA-seq using wildtype (Prp2) RNA. **(B)** A scatter plot depicts intron-retention measurements in MPE-seq and conventional RNA-seq using RNA from a splicing mutant strain (*prp2-1*). **(C)** A scatter plot depicts the fold-change (*prp2-1*/Prp2) in intron-retention as measured by MPE-seq and conventional RNA-seq

**FigS4:**
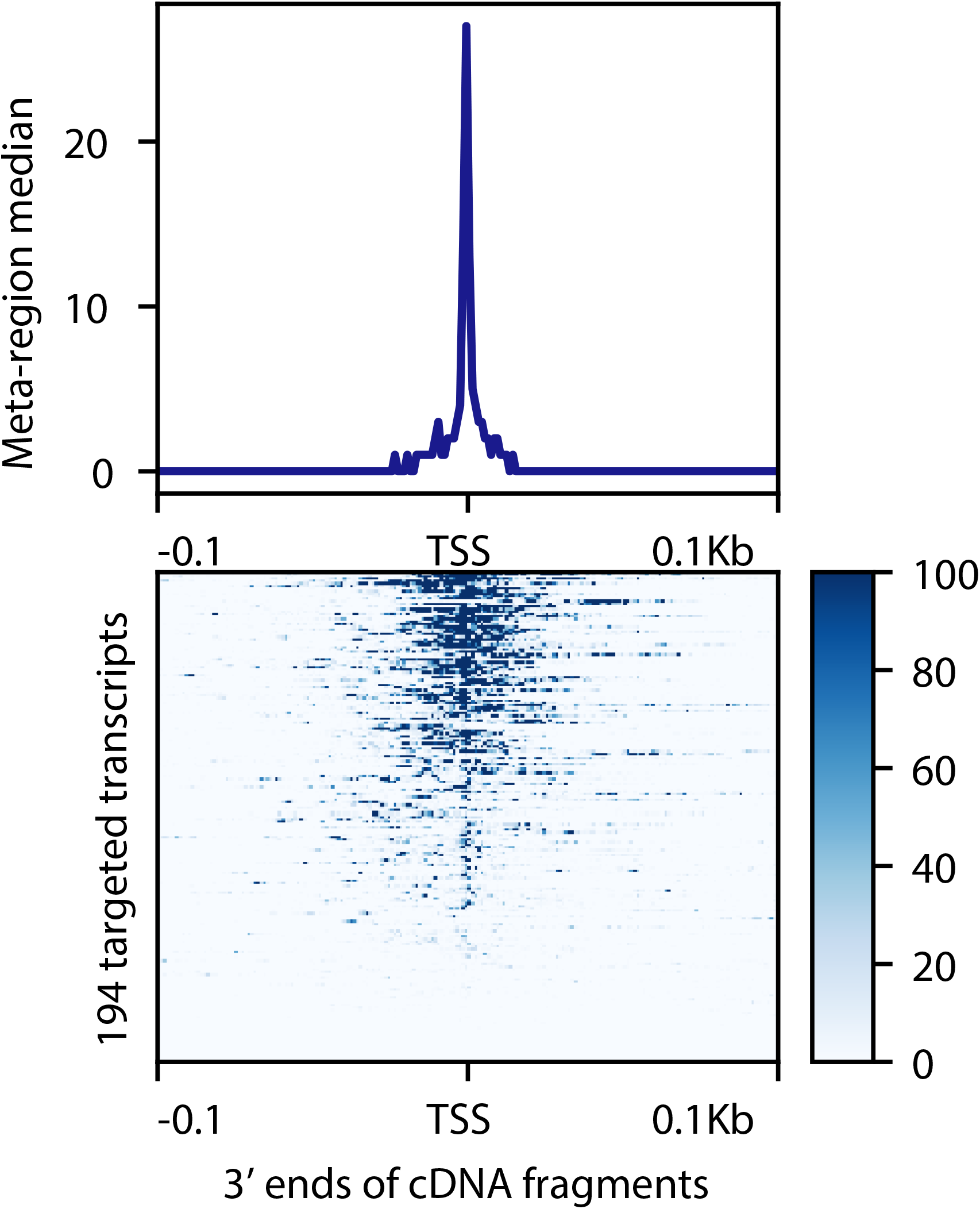
Transcription start site profiling by MPE-seq. Metagene profile of 3’ ends mapped by MPE-seq, centered around transcription start sites (TSS) as determined by PRO-cap, an orthologous method for mapping transcription start sites. The high abudance of read ends that pile up at TSSs indicates that MPE-seq can be used to profile cDNA termini.

**FigS5:**
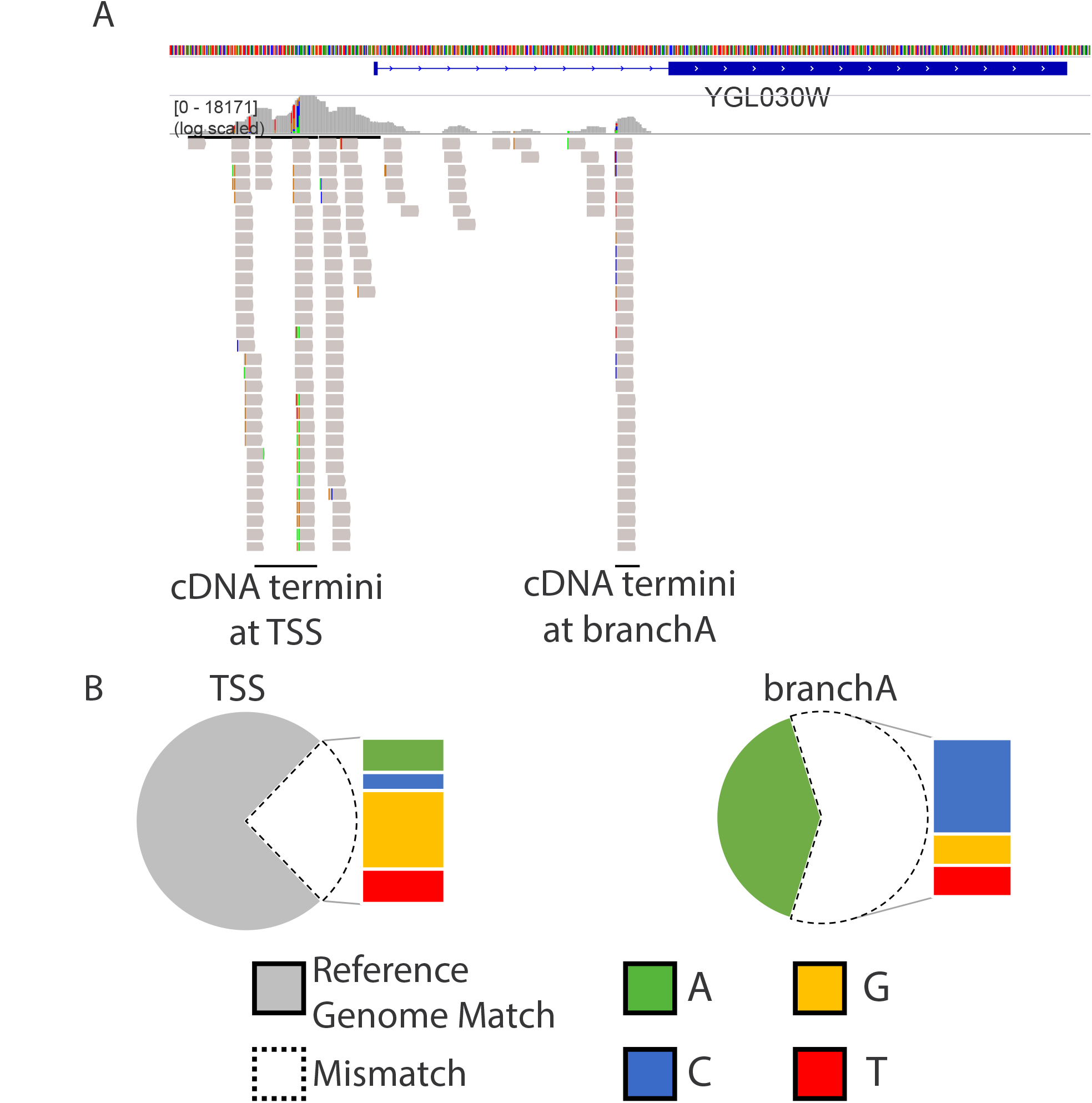
Lariat-intermediate derived cDNAs contain a unique signature of mismatches incorporated by reverse transcriptase at the branched adenosine (A) Genome browser screenshot of 3’ reads from paired-end sequenced fragments illustrates the unique signature of non-templated base incorporation by reverse transcriptase at a branched-adenosine vs the 5’ RNA terminus (B) Genome-wide quantification of the mismatch frequencies at 3’ cDNA termini near the TSS (left) versus at the annotated branchpoint (right).

**FigS6:**
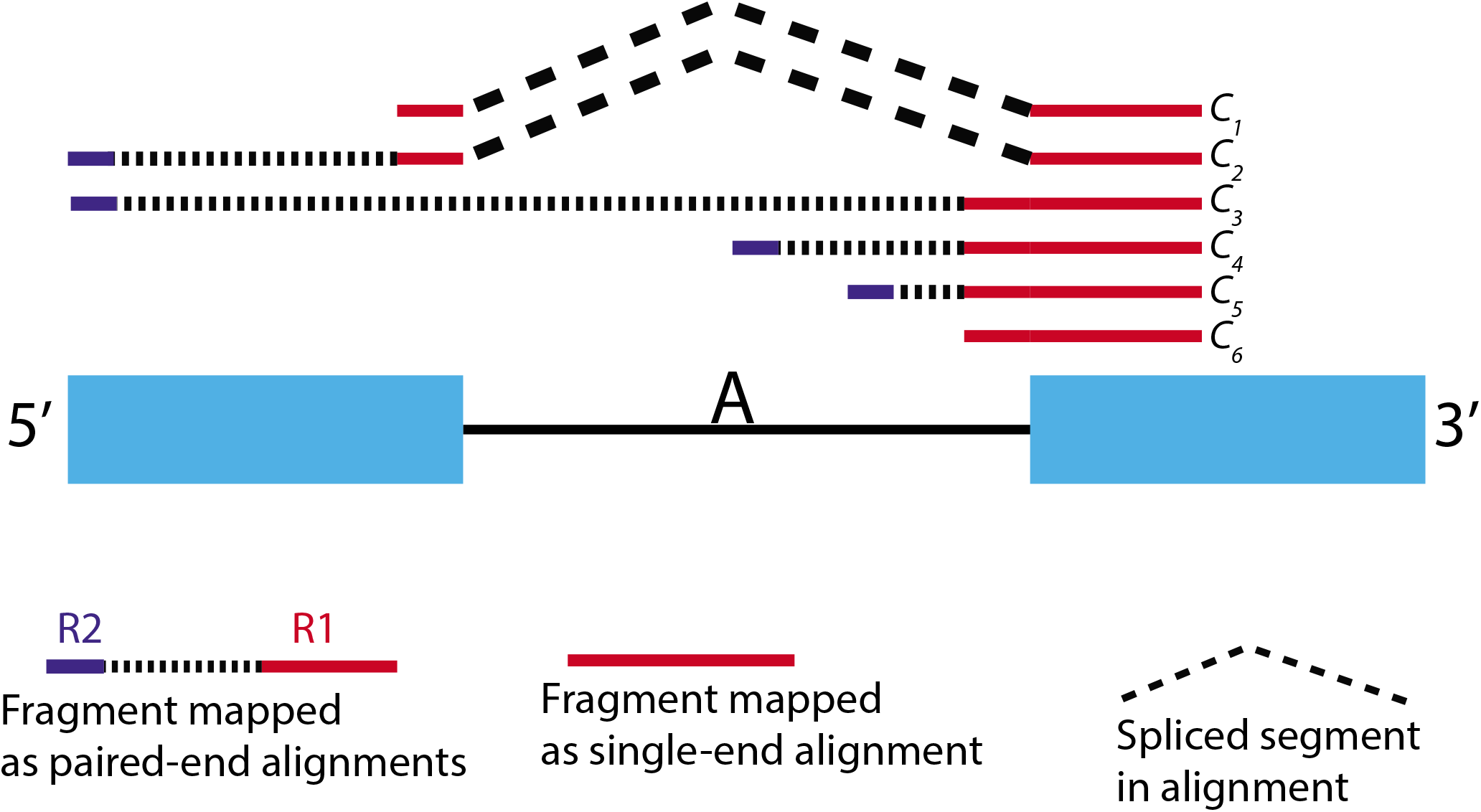
Schematic for assigning reads to splice intermediate isoforms. To quantify the abundance of pre 1^st^ step, lariat intermediate, and spliced isoforms for each targeted splice event, we categorized fragments into six classes based on paired-end read alignments. Fragments containing a splice junction (C_1_ and C_2_) are indicative of a spliced RNA (S). Fragments that are unspliced and traverse the branchpoint region (C_3_) are classified as pre-1^st^ step RNA (P). Fragments that are unspliced but terminate within a −3 to +5bp window from the previously determined branchpoint (C_4_) are classified as lariat intermediate (L). Fragments that are unspliced and either terminate downstream of the branchpoint (C_5_) or the terminus could not be mapped (C_6_) are ambiguous between P and L. Therefore, for accounting purposes, the counts for these fragments were coerced into P and L classifications based on the ratio of P and L determined by unambiguous mappings (C_3_ and C_4_). See methods for more details.

**FigS7:**
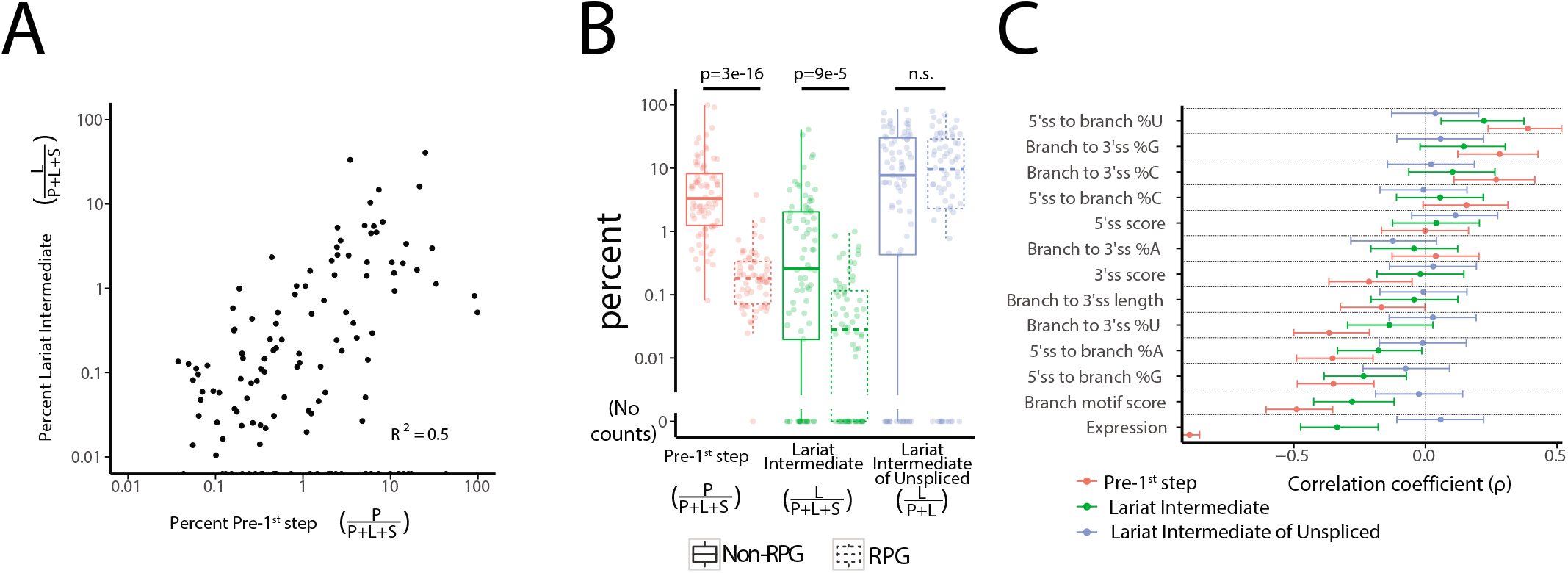
Transcript features that correlate with the abundance of lariat intermediate. (A) Scatter plot depicting the correlation between the abundance of pre-1^st^ step RNA and the abundance of lariat intermediate (B) The abundance of pre-1^st^ step RNA and lariat intermediate RNA is significantly correlated with classification of introns into those that are in ribosomal protein genes (RPG) and non-RPG. However, the abundance of lariat intermediate relative to pre-1^st^ step RNA, a metric of the efficiency of the second step of splicing, does not correlate. **(C)** Spearman correlations of various features to the abundance of pre-1^st^ step RNA, lariat intermediate, the abundance of lariat intermediate relative to pre-1^st^ step RNA. Errorbars indicate 95% confidence intervals as estimated by Fisher transformation of Spearman’s correlation coefficient.

**FigS8:**
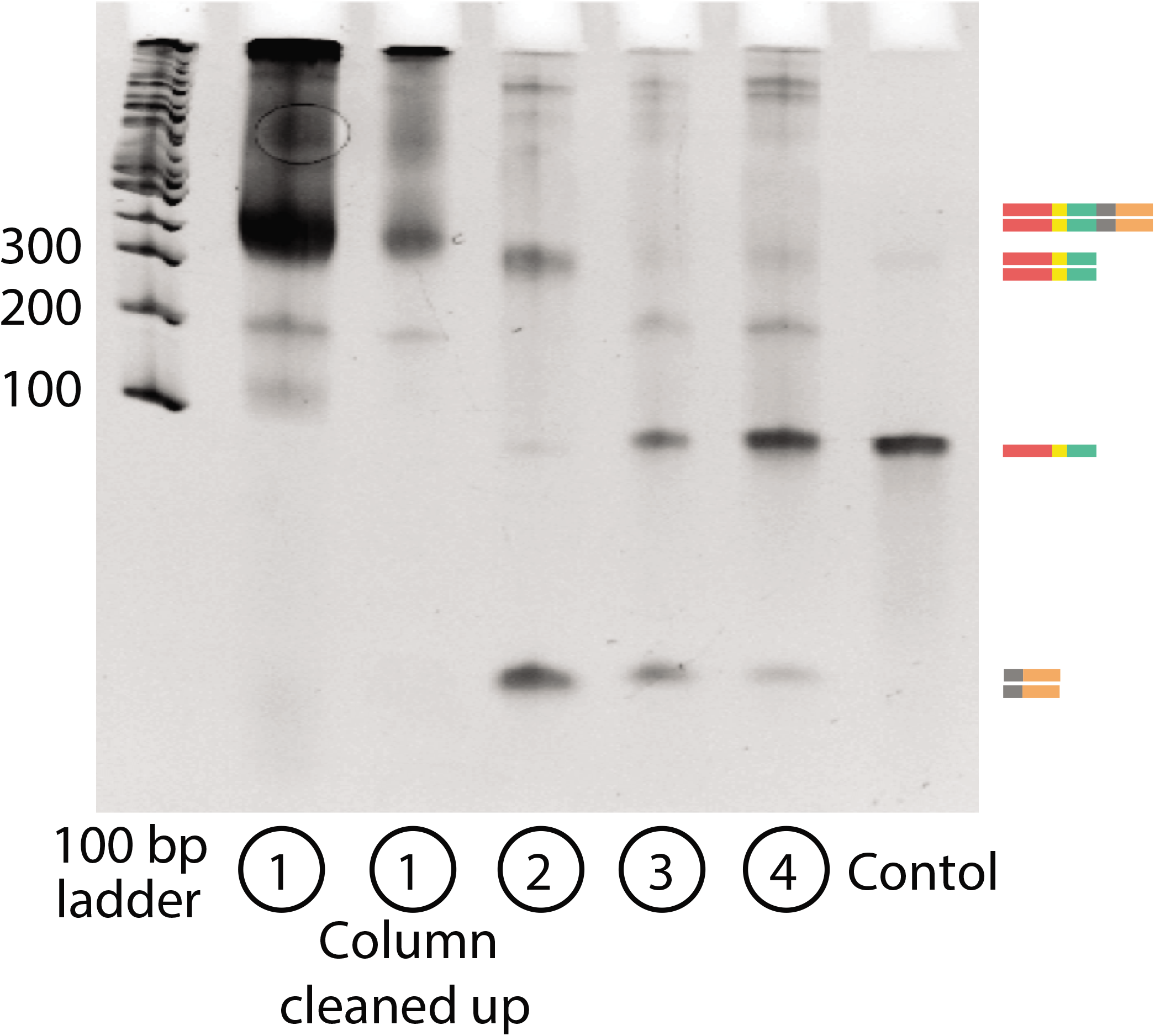
Array-based oligonucleotide synthesis can be used to generate primer pools for use in MPE-seq. Steps during the amplification and purification of array-synthesized primer pools are monitored via native gel electrophoresis. The control lane represents a pool of individually synthesized MPE-seq primers which did not require amplification and purification. Lanes refer to products of each individual step in the protocol. (1) PCR amplification of the oligonucleotide synthesis pool using a 5’ blocked sense primer and a biotinylated antisense primer. (2) Restriction digestion to cleave off the PCR primer handle. (3) Lambda exonuclease digestion of free 5’ ends. (4) Streptavidin purification of biotinylated PCR handle. The unbound fraction is the desired primer pool product.

